# Diverse digital and fuzzy composite transcriptional elements are prevalent features of mammalian cis-regulomes

**DOI:** 10.1101/2021.11.26.470154

**Authors:** Virendra K. Chaudhri, Harinder Singh

**Affiliations:** Center for Systems Immunology and Department of Immunology, University of Pittsburgh, Pittsburgh, Pennsylvania 15213; Department of Computational and Systems Biology, University of Pittsburgh, Pittsburgh, Pennsylvania 15213

## Abstract

Mammalian transcriptional regulatory sequences are comprised of complex combinations of simple transcription factor (TF) motifs. Stereospecific juxta-positioning of simple TF motifs generates composite elements (CEs), that increase combinatorial and regulatory specificity of TF-DNA interactions. Although a small number of CEs and their cooperative or anti-cooperative modes of TF binding have been thoroughly characterized, a systematic analysis of CE diversity, prevalence and properties in cis-regulomes has not been undertaken. We developed a computational pipeline termed CEseek to discover >20,000 CEs in open chromatin regions of diverse immune cells and validated many using CAP-SELEX, ChIP-Seq and STARR-seq datasets. Strikingly, the CEs manifested a bimodal distribution of configurations, termed *digital* and *fuzzy*, based on their stringent or relaxed stereospecific constraints, respectively. *Digital* CEs mediate cooperative as well as anti-cooperative binding of structurally diverse TFs that likely reflect AND/OR genomic logic gates. In contrast, *fuzzy* CEs encompass a less diverse set of TF motif pairs that are selectively enriched in p300 associated, multi-genic enhancers. The annotated CEs greatly expand the regulatory DNA motif lexicon and the universe of TF-TF interactions that underlie combinatorial logic of gene regulation.

---

DNA-protein interactions mediated by combinations of transcription factors (TFs) are central to the decoding of regulatory information in metazoan genomes(*1–5*). Most eukaryotic TFs bind to simple DNA motifs (6-10 bp) that display significant redundancy(*3*). As a consequence, simple TF motifs have low information content and insufficient regulatory potential on their own. Promoters and enhancers of metazoan genes are comprised of complex combinations of simple TF motifs(*2, 4*). Despite numerous attempts to unravel the structural and functional basis of these higher order TF-motif configurations, the nature of the code used for storing and retrieving transcriptional regulatory information has remained elusive. A small set of TF motifs have been shown to form composite elements (CEs) that promote the cooperative binding of cognate TFs belonging to structurally distinct families(*6–11*). Exemplars described by us and others include the EICE motif recognized by Ets (PU.1, SpiB) and IRF (IRF4, IRF8) family members, the NFAT-AP1 (JunB:Batf) motif bound by NFAT and IRF family members and the AICE motif recognized by AP1 and IRF (IRF4, IRF8) family members(*12–14*).

Composite transcriptional elements are typically constrained by the relative orientation and spacing of their constituent simple motifs. CEs have higher information content, that can be computed on the basis of the two parent motifs comprising them as well as the length of the spacer (see below), and consequently generate greater regulatory specificity. The diversity and prevalence of CEs within eukaryotic regulatory DNA sequences remain to be systematically delineated. Earlier efforts to extensively analyze CEs have been constrained by their focus on limited genomic search space or biochemical screens involving synthetic DNA libraries of binary TF motifs(*15–19*). Recently extensive genomic analyses with a multitude of cell types and states have generated comprehensive datasets that facilitate a systematic search for CEs (*20*). We reasoned that a systematic analysis of such datasets could address a long-standing question of whether CEs are diverse and widely occurring in genomic regulatory sequences. If so, then these binary motifs would greatly expand the lexicon of the genomic regulatory code and in turn the catalog of TF-TF interactions that underpin combinatorial control of gene regulation.

The ImmGen consortium has systematically profiled, using standardized work flows, the chromatin structures of 86 hematopoietic and immune cell types and their varied developmental or activation states using ATAC-seq(*20*). This unparalleled and rigorously curated catalog of open chromatin regions (OCRs) has delineated >500,000 presumptive transcriptional regulatory sequences that function to regulate gene activity in diverse cellular contexts. We reasoned that this exceptional resource would enable a comprehensive discovery of CEs and a determination of their prevalence as well as structural diversity. In our computational framework, we allowed for variable spacing of the two TF binding sites or motifs that constitute a CE. For any pair of TF motifs separated by a given distance (base pairs, bp), there are four possible configurations (Figure 1A). We developed an R based pipeline(*21*), CEseek, that searches a test set of regulatory sequences along with a background set for all possible CE configurations, with variable spacing, for pairs of TF motifs each of which are described by position probability matrices (PPMs) (Figure 1B, see Methods). Sequences of CEs that are statistically enriched in the test set in relation to the background set are then used to generate an output file of CEs along with their PPMs. To validate the CEseek pipeline, we scanned a set of murine regulatory sequences to determine if it could recover previously characterized CEs (EICE, AICE and NFAT-IRF motifs). For this application, a set of functionally tested B cell enhancer sequences(*22*) along with a 10-fold scrambled set of background sequences were scanned for CEs comprised by the PU.1, IRF, AP1, and NFAT motifs. The analytical method recovered all three CEs, notably both configurations of AICE (0 and +4bp spacing) (Figure 1C), that have previously been characterized thoroughly using biochemical, functional and genomic *assays*(*12–14*).

**Fig. 1.**
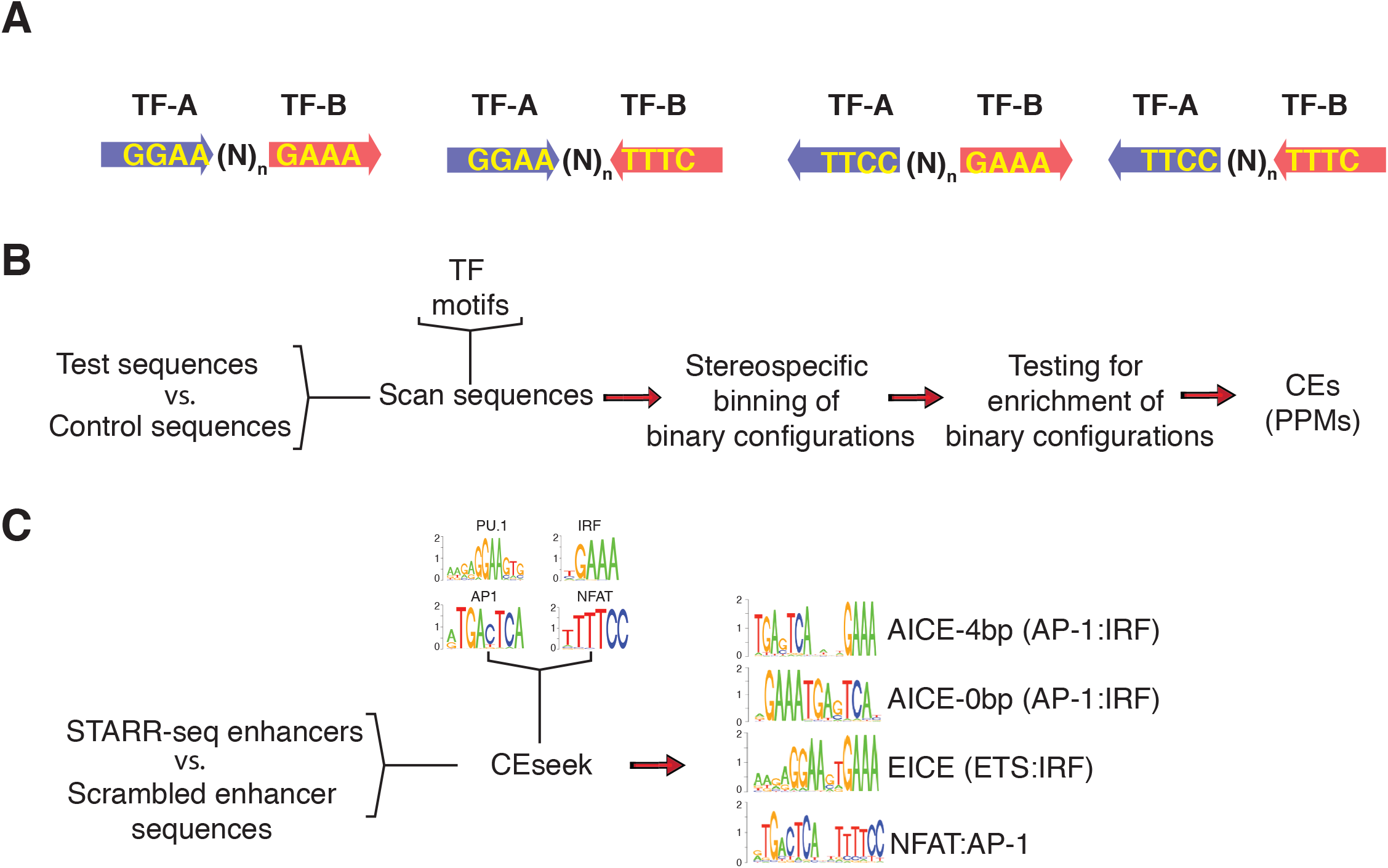
CEseek, a computational pipeline for discovery of Composite Elements in regulatory DNA sequences. (A) Schematic illustrating possible configurations of a CE constituted by stereo-specific pairing of non-palindromic DNA binding motifs (TF-A and TF-B) and allowing for variable spacer lengths. (B) Computational work flow for CEseek using test and control sequences. (C) CEseek recovered known composite regulatory elements comprised by motifs for the transcription factors PU.1, IRFs, AP1 family members and NFATs in a set of B cell enhancers (~12,000) delineated by STARR-seq(*22*). Motif logos for AICE, EICE and AP1:IRF BiTs are displayed (logP values: AICE(4bp) = −26.87, AICE(0bp) = −15.05, EICE = −149.75, NFAT:AP1 = −5.85).

To systematically analyze the identities and occurrence of CEs in a large set of presumptive transcriptional regulatory sequences, we performed an all-by-all TF motifs scan of the ImmGen ATAC-seq dataset(*20*). As noted above, this database catalogs the open chromatin regions (OCRs) of a total of 86 immune cell types and states along with 4 non-hematopoietic cell types and is comprised of 512,292 OCRs, each 180bp in length (For details see Methods). The sequences were scanned by CEseek, using a consolidated set of 153 core TF motifs compiled from CisBP and HOMER (*23, 24*)(Supplementary Table 1, see Methods) that were representative of structurally diverse families of mammalian TFs. The total number of binary TF motif combinations, termed CE types, used to scan the 512,292 ImmGen OCRs with CEseek was n=11,628. Scanning for CEs was performed using an end-to-end separation distance of −5 to 10 bp for the individual TF motifs (Figure 2A). We allowed for partial overlap of the individual motifs based on three considerations (i) the base pairs flanking core TF motif residues have lower information content(*25*) (ii) cooperative binding can be associated with alterations in DNA binding specificity of one or both of the TFs(*19*) and (iii) partially overlapping motifs can lead to anti-cooperative binding by the two TFs(*26, 27*). Allowing for these flexible spacing (16 spacer lengths) and orientation possibilities (4), the 153 core TF motif combinations yielded 744,192 (n=11,628×16×4) possible CE configurations. We note that CEseek allows the user to specify spacer length constraints, in our analysis each CE type could manifest 64 (n=16×4) possible configurations. After the initial CEseek scan that resulted in binning of all possible CE hits, the prevalence of each of these configurations was tested for their enrichment in cell-type specific OCRs. We reasoned that selective enrichment of a subset of all possible CEs in cell-type specific OCRs would not only provide statistical validation but also biological support for their functional importance. To enable this analysis, each ImmGen cell type/state was associated with OCRs that were preferentially accessible in its chromatin landscape in relation to the large majority of the other cell types/states. Accordingly, we annotated each ImmGen cell type/state with a distinctive set of OCRs that manifested ATAC-seq peak signal intensities that were within the 95^th^ percentile (most-accessible) based on analysis of the distributions of individual peak signal intensities across all 90 cell types/states. This enabled developmentally regulated OCRs to be associated with more than one cellular state within a given differentiation trajectory. Enrichment of each possible CE configuration was then tested by using the aggregated set of cell-type/state specific OCRs as a test set and all remaining OCRs in the ImmGen dataset as the background set. CEs that were statistically enriched in cell type/state-specific OCRs were delineated using a thresholding p-value of less than 10^-20^, informed by the total numbers of sequences in the test and background sets. This analytical process resulted in the identification of 22,698 CEs that were dominant DNA sequence features of immune cell type/state-specific OCRs and are termed ImmCEs (Figure 2A, Supplementary Table 2). The entire set of ImmCE configurations (n=22,698) were represented by 2,992 distinct TF motif pairs (ImmCE types) within a possible set of 11,628 TF motif pairs used in the CEseek scan. Notably, the EICE, AICE and NFAT-AP1 composite elements were recovered within the very large set of ImmCEs. As expected, ImmCEs had larger sequence lengths (bp) compared with their simple parent motifs (Figure 2B). Exemplary ImmCEs comprised of Runx1-Prdm14 and Stat5-RBPJ (Notch) motifs are shown (Figure 2C). The vast majority of ImmCEs were different in sequence information content than the parent motifs from which they were derived. Even allowing for significant overlap between the simple TF motifs (−5 to −1 bp), only 552 of the 22,698 ImmCEs showed a PPM similarity score of 0.90 or greater to one of the parent motifs (Supplementary Table 3). Since the ImmCEs were delineated based on their statistical enrichment in cell type/state-specific OCRs using the remaining ImmGen OCRs as a background set, we independently tested for the prevalence of these CEs in the entire set of ImmGen OCRs using a complementary approach, that utilized a scrambled set (5x) of ImmGen OCR sequences as the background. This complementary statistical analysis revealed 16,839 of the ImmCEs to be significantly enriched in the test set with p values of less than 10^-10^ (Supplementary Table 4). As a control, a similar enrichment analysis using all ImmGen OCRs and their scrambled counterpart sequences as background was performed with the parent 153 TF simple motifs. The comparative analysis showed ImmCEs to be enriched much more than the simple motifs from which they were derived, with the top quartile having fold-enrichment values of greater than 16 in ImmGen OCRs. (Fig 2D). Given the enrichment of ImmCEs in cell type/state-specific OCRs we hypothesized that they could be used to predict, without any additional priors, the developmental connectedness of immune and hematopoietic cells based on their prevalence in context-specific regulatory genomic landscapes. To test the hypothesis we constructed a minimal spanning tree (MST) using as features the frequencies of occurrence of the most highly enriched ImmCEs (top 2 quartiles of distribution in Fig. 2D) within cell type/state-specific OCRs. The MST generated 5 major clusters that segregated distinct groups of developmental intermediates and mature immune cells (Figure 2E). Notably, hematopoietic stem cells, their multipotential progenitors and early B- and T-lineage progenitors formed one major branch. A second cluster comprised of natural killer (NK) cells and innate lymphocytes (ILCs). Mature and activated B and T cells were largely separated into distinct clusters. Finally, monocytic and dendritic cells represented a distinctive major cluster. These major as well as minor clusters are consistent with the known developmental relationships of various immune progenitors and their mature progeny(*20*). It is important to note that this analysis solely utilized ImmCEs with high enrichment values and their prevalence in cell type/state-specific OCRs with no knowledge of the underlying transcriptional landscapes. Thus, the ImmCEs appear to represent key elements of the regulatory genomic codes that are used in specifying immune cell identities.

**Fig. 2.**
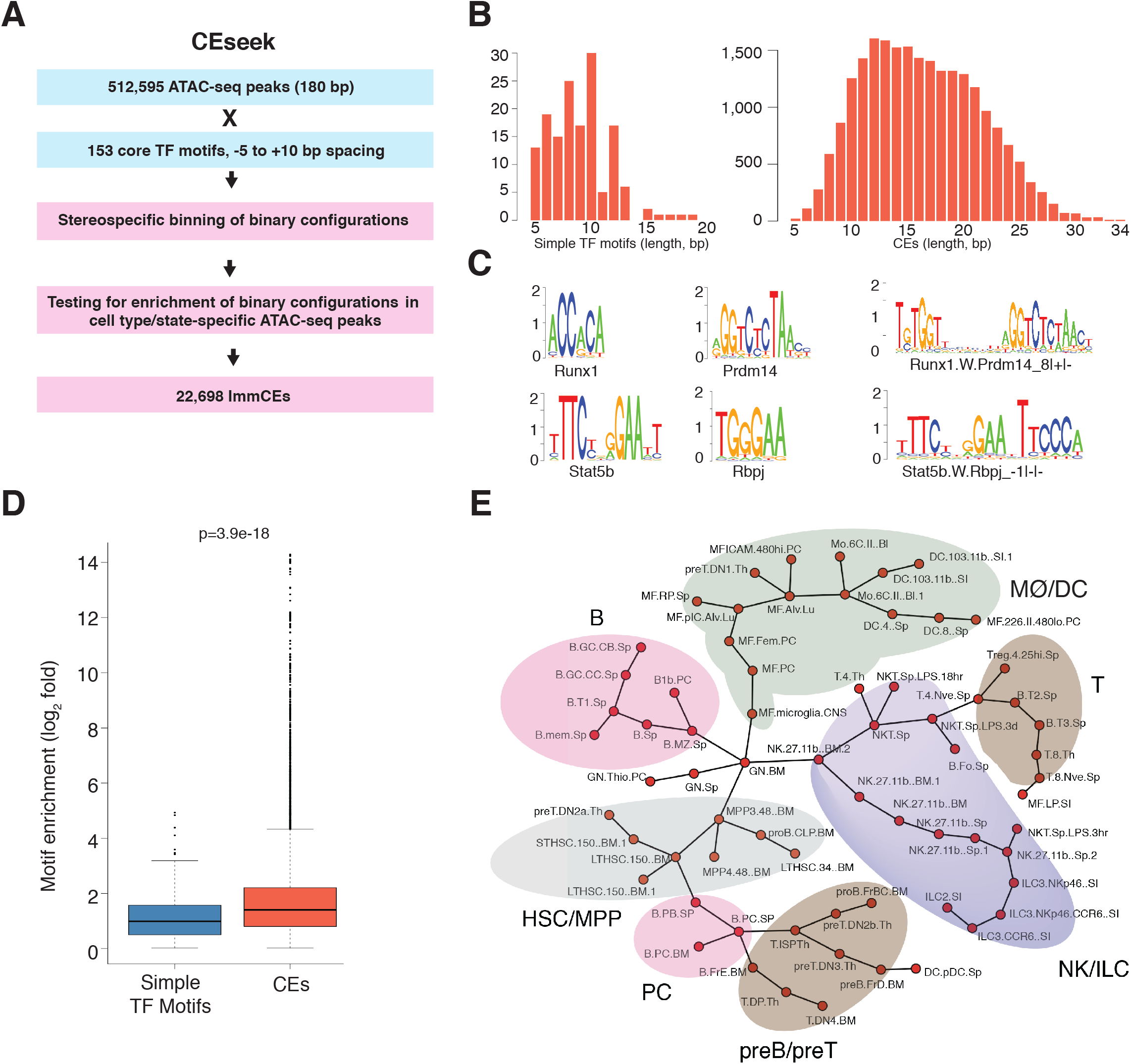
Discovery and analysis of CEs in regulatory DNA sequences of diverse immune cell types and differentiation states. (A) Computational workflow for discovery of CEs in ImmGen ATAC-seq datasets encompassing 90 distinctive cell states(*20*). (B) Length distributions of simple TF motifs versus CEs delineated by CEseek. (C) Two exemplary CEs constituted by juxta positioning of the indicated TF motifs (D) Enrichment of CEs versus simple TF motifs in ImmGen ATAC-seq datasets. Background set was generated by scrambling test sequences 5-fold. Log2 fold enrichment values for CEs with logP < −10 are plotted. (E) Minimal spanning tree analysis of the relatedness of immune cell types and their differentiation states based on the frequency of occurrence of specific CEs (logP < −10) in cell type-specific OCRs.

We next analyzed the diversity of TF motif pairs (CE types) as a function of the number of possible configurations for each motif pair (Figure 3A). Strikingly, the largest number of TF motif ImmCE types were defined by only one of 64 possible configurations. In fact, the vast majority of ImmCE types (2,418 of 2,992) were delineated by 6 or less of 64 possible configurations. These ImmCEs display stereospecific constraints in the juxta-positioning of the TF motifs comprising them. Hereafter, these will be termed *digital* CEs (Supplementary Table 5). The distribution of ImmCE types as a function of numbers of permissible configurations had an unexpectedly long tail that extended to all possible 64 configurations for a small number of CE types. This unusual feature was readily apparent when the TF motif pair counts (Figure 3A) were transformed by multiplying each with their corresponding numbers of observed configurations. This resulted in a bimodal CE count distribution that reflected ImmCE types with sparse or numerous configurations (Figure 3B). The left end of the distribution (CE types with 6 or less configurations, n=2,418 representing 5,063 configurations) was reflected by the *digital* CEs (exemplars shown in Figure 3C) whereas the right end of the distribution (CE types with 49 or more possible configurations, n=170 representing 9,399 configurations) was reflected by TF motif pairs lacking stereospecific constraints and therefore termed *fuzzy* CEs (exemplars shown in Figure 3D). Given the lack of spacing constraints manifested by *fuzzy* CEs, they were longer in overall length (median, 17 bp) compared with *digital* CEs (median, 13 bp) (Figure 3E, F). We next analyzed the information content of *digital* and *fuzzy* CEs using motif scores (Biostrings maxScore, Figure 3G). In this approach, the PPM of the entire CE, comprised by the two TF motifs along with the spacer, is used in computing the overall motif score. Longer spacer lengths result in higher motif scores. Thus, given their overall longer length distribution, *fuzzy* CEs had a higher median motif score compared with *digital* CEs, but both reflected significantly greater sequence complexity than the simple motifs from which they were derived. Detailed analysis of the spacing distributions of *digital* and *fuzzy* CEs revealed that although they spanned the entire spectrum of −5 to +10 bp, as expected they were oppositely shifted in terms of spacer lengths (Supplementary Figures S1 and S2). We note that partially overlapping CE configurations predominated in the *digital* distribution but in spite of their inter-digitation manifested discernable pairings of their constituent simple motifs (Supplementary Figure S1). To gain insight into the types of TFs that could recognize *digital* and *fuzzy* CEs we first rank ordered TF motifs on the basis of their extent of occurrence in distinct CE types as well as on their occurrences in the total counts of CE configurations (Supplementary Figures S3, S4). The former distribution highlighted motifs for TFs such as NFe2l1, Rara, Nfic (Supplementary Figures S3A, S4A,B) whereas the latter featured E2f1, Zbt7b and Klf7 (Supplementary Figures S3B, S4C,D). These results suggest that TFs such as NFe2l1, Rara, Nfic appear to interact with a diverse set of cooperative or antagonistic TFs but in stereospecific configurations. In contrast, the TFs such E2f1, Klf7 and Zeb1 are inferred to preferentially engage with restricted sets of partner TFs on their cognate CEs, via mechanisms that are not dependent on stereospecific DNA binding (discussed below).

**Fig. 3.**
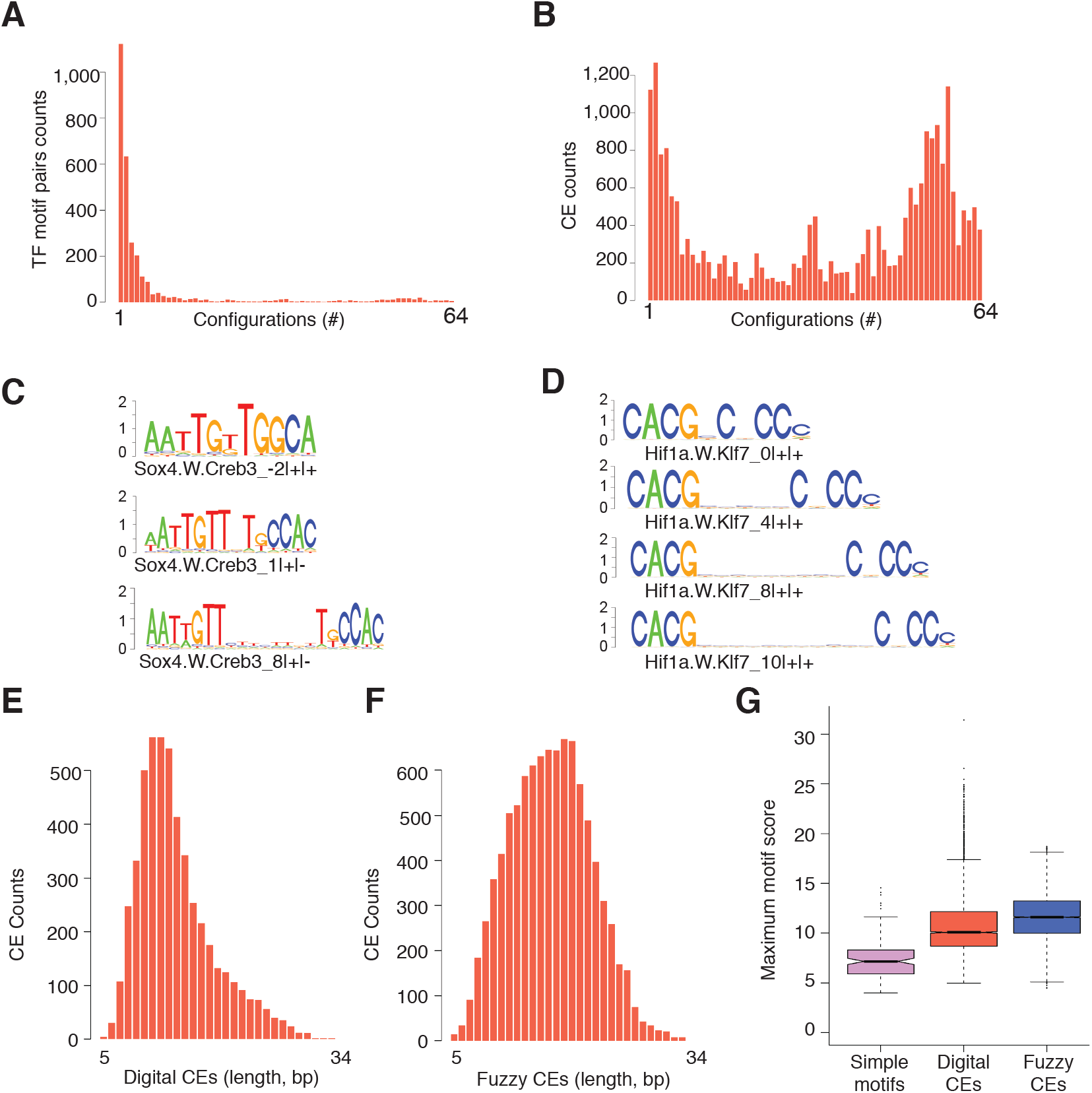
CEs comprise *digital* as well as *fuzzy* configurations. (A) Diversity of TF motif pairs (CE types) comprising digital (up to 6 configurations) and fuzzy CEs (> 48 configurations). (B) Frequency distribution of digital and fuzzy CE configurations. (C) Representative digital CEs for Creb3 and Sox4 TF motifs are shown as motif logos for 3 configurations. Notations below each CE motif logo indicate the paired (W) simple motifs with their spacer configuration denoted as −5 to +10 followed by the orientations (+/-) of the parent TF motifs. (D) Representative fuzzy CEs for Klf7 and Hif1α TF motifs are shown as motif logos for 4 out of 51 configurations (C) Length distribution plot of digital CEs. (D) Length distribution plot of fuzzy CEs. (E) Box plots showing information content distributions of simple motifs, digital CEs and fuzzy CEs based on maximum scores of their constituent motifs. Notches indicate 95% confidence intervals.

Each ImmCE in our regulatory DNA catalog is annotated for the OCRs in the ImmGen datasets within which it resides. Each such CE in turn generates a multitude of molecular, genomic and functional predictions. We reasoned that we could systematically test for the molecular and biological significance of a large number of ImmCEs given the availability of informative datasets spanning TF-TF cooperative DNA binding, TF-TF co-occupancy in genomic regions and functionally validated enhancers. As a first step we tested if the predicted *digital* CEs likely promote cooperative DNA binding by cognate TFs belonging to structurally distinct families. To do so, we took advantage of a unique CAP-SELEX dataset(*19*). This dataset was generated by analyzing cooperative binding of a large number of TF pairs to synthetic DNA sequences *in vitro* using recombinant proteins and consecutive affinity-purification systematic evolution of ligands by exponential enrichment (CAP-SELEX) analysis. The biochemical screen identified 315 TF-TF pairs that could recognize 618 composite elements with characteristic orientation and spacing of their simple motifs. We reasoned that if such TF-TF cooperative binding interactions are biochemically and biologically relevant then many of the composite elements revealed by CAP-SELEX should be contained within our larger set of *digital* CEs (n=5,063) which are conserved features within a large set of genomic regulatory sequences. We therefore compared the CEs within the CAP-SELEX and CEseek datasets based on their PPMs. Majority of the CAP-SELEX CEs (75%) had similarity scores of greater than 0.7 when compared with CEseek CEs (Figure 4A). Representative CAP-SELEX CEs that are highly related to CEseek CEs are shown in Figure 4B. Remarkably, these matches based on similarity scores ranging from 0.79 to 0.93 revealed strong alignments with CEseek predicted CEs that displayed either gapped or overlapping TF motifs. We note that CAP-SELEX CEs were based on successive affinity selection of DNA sequences using in vitro DNA binding assays and thus are more constrained by a focus on highest affinity sequences whereas CEseek delineated CEs represent naturally varying endogenous regulatory motifs. Furthermore, CEs delineated by CEseek using the ImmGen dataset are reflective of regulatory motifs enriched in cis-regulomes of immune cells whereas the CAP-SELEX screen imposed no biological context or constraint. In spite of these crucial differences between the two methodologies, the majority of the CAP-SELEX CEs were recovered within the larger set of *digital* ImmCEs. Strikingly, the distribution of TF-TF pair counts as a function of numbers of configurations in CAP-SELEX CEs (Supplementary Figure S5A) was very similar to that displayed by *digital* CEseek CEs (Figure 3A). Given that ImmCEs not only contain a very large set of novel CEs but also new configurations of previously characterized CEs, we tested examples of both types by biochemically analyzing a novel NFAT:IRF motif and new configurations of the well characterized EICE motif (see Extended Data Figures). The former promoted cooperative binding by NFATc2 and IRF4 or IRF8 whereas the latter EICE configurations, in spite of their varying spacer lengths (1, 3 and 4 bp), sustained cooperative binding by PU.1 and IRF4 or IRF8. Importantly, the novel NFAT:IRF CE as well as new configurations of EICE that were cooperatively bound by their cognate TFs also promoted transcription in a reporter assay. Thus, the larger set of *digital* CEs delineated by CEseek are strongly predictive of cooperative binding by pairs of TFs and greatly expand the universe of such TF-TF interactions.

**Fig. 4.**
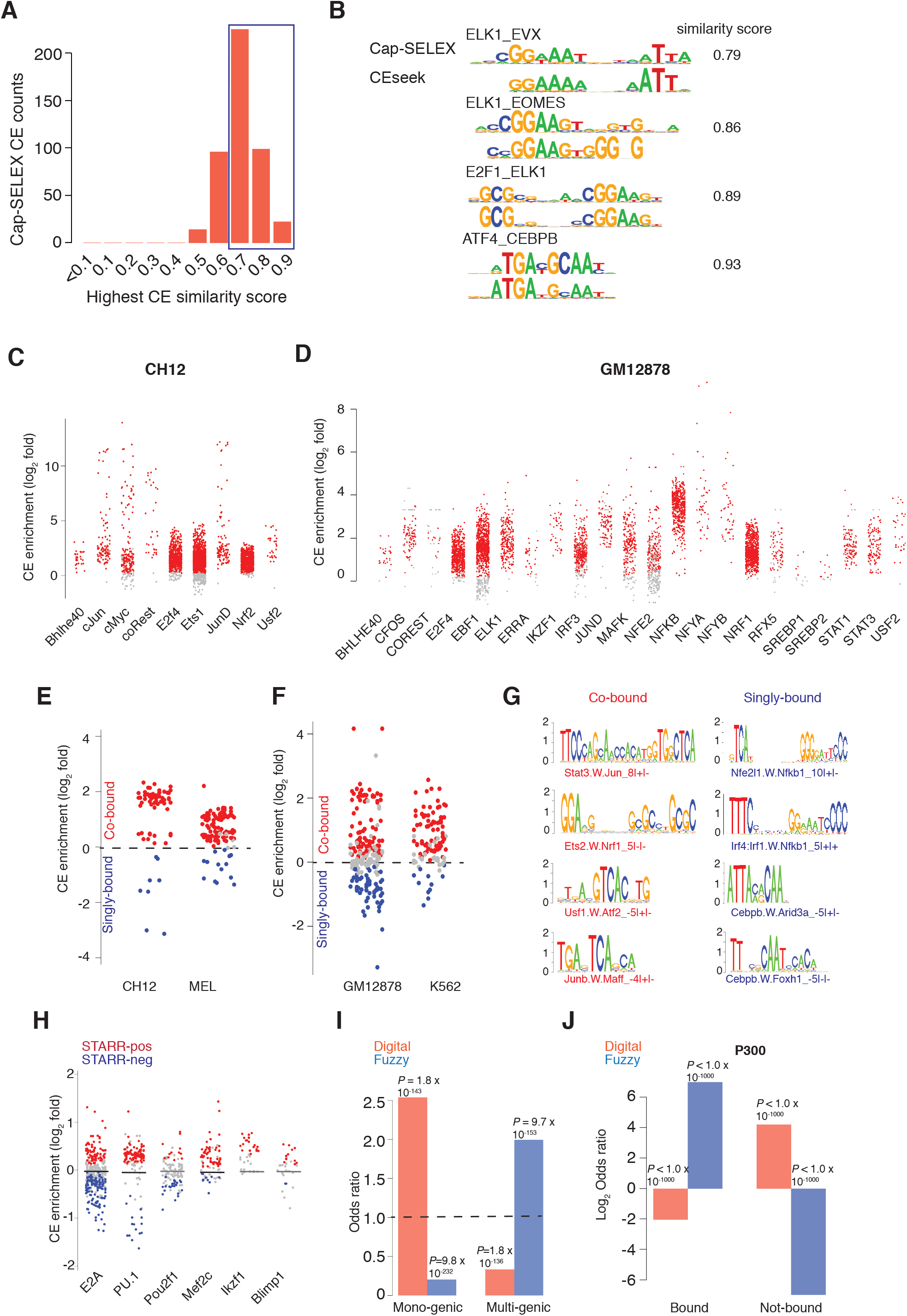
Biochemical, genomic and functional validation of CEs. (A) Composite TF motifs identified by in vitro DNA binding assays (Cap-SELEX)(*19*) are contained within CEs revealed by CEseek. The distribution shows the highest similarity scores of Cap-SELEX motifs with cognate CEs. (B) Representative Cap-SELEX motifs aligned with cognate CEs delineated by CEseek, along with their motif similarity scores. (C) Enrichment of CEs for the indicated murine TFs in their ChIP-seq peaks in the murine CH12 B cell line(*28*). For this analysis, a background set of sequences was constituted by scrambling the test set (10x). Red dots indicate CEs with an enrichment significance of logP < −20. Grey dots are CEs that do not satisfy the significance criterion. (D) Enrichment of CEs for the indicated human TFs in their ChIP-seq peaks in the human GM12878 B cell line(*28*). Background set of sequences was generated as in panel C. (E) Analysis of co-bound and singly-bound genomic regions, containing predicted CEs, by their cognate TFs in indicated murine cell lines. Red dots represent CEs that are statistically enriched (logP < −10) in co-bound regions of relevant pairs of TFs in their ChIP-seq datasets. Blue dots represent BiTs that are statistically enriched (logP < −10) in singly bound regions of relevant pairs of TFs. Grey dots represent BiTs that are not significantly enriched in either co-bound or singly bound regions. (F) Analysis of co-bound and singly-bound genomic regions, containing predicted CEs, by their cognate TFs in indicated murine cell lines (colors of dots are as in panel E). (G) Representative CEs that appear to promote cooperative (red labels) or anti-cooperative (blue labels) TF binding *in vivo* are displayed with their motif logos. (H) Enrichment of CEs for the indicated TFs in active B cell enhancers identified by STARR-seq(*22*). Red or blue colored dots represent logP values < −5 for enrichment in STARR-pos(n=11,809) and STARR-neg (n=11,942) regions respectively (I) Differential enrichment of digital and fuzzy CEs in mono-genic (n=649) and multi-genic (n=2,263) B cell enhancers(*22*). Odds ratios are plotted for the counts of digital or fuzzy CEs enriched with a logP < −5. (J) Differential enrichment of digital and fuzzy CEs in STARR-seq B cell enhancers bound by P300 (n=5,511) vs rest of STARR-seq B cell enhancers (n=6,298)(*22*). Log_2_ values of odds scores for counts of enriched digital and fuzzy CEs with a logP < −5 are plotted.

Next to test the genomic binding patterns of cognate TFs predicted by the CEs *in vivo*, we reasoned that analyses of their distributions in relevant TF ChIP-seq datasets(*28*) would not only enable us to determine their enrichment in predicted TF binding regions but also to distinguish between cooperative or anti-cooperative DNA binding modes by the cognate TF pairs. The latter aspect would be reflected by their co-or mutually exclusive genomic occupancy at such regions, respectively. We first tested for enrichment of cognate CEs using ENCODE ChIP-seq datasets for their corresponding TFs in the murine hematopoietic cell lines, CH12 (B-lineage) and MEL (erythroid). Majority of the predicted CEs for the indicated TFs were highly enriched in their ChIP-seq peak regions in both cellular contexts (red dots) (Figure 4C and Supplementary Figure S5B). Given that ENCODE has profiled a larger set of more diverse TFs in human cell lines and that we wanted to determine if the murine CEs were conserved in human cis-regulatory sequences we analyzed for CE enrichment in ChIP-seq data sets for the ENCODE cell lines GM12878 (B-lymphoblastoid) and K562 (erythromyeloid). As was the case above, a large fraction of cognate CEs for the predicted TFs were enriched in their genomic DNA binding regions (Figure 4D and Supplementary Figure S5C). Given that the majority of the CEs that were tested for their enrichment in cognate TF ChIP-seq datasets were *digital* in nature, we performed a focused analysis of *fuzzy* CEs with their cognate TFs. Although *fuzzy* CEs involved a less diverse set of TFs they like their *digital* counterparts were significantly enriched in murine and human ChIP-seq datasets (Supplementary Figures S5D-G).

The availability of ENCODE TF ChIP-seq datasets for pairs of TFs, for which CEseek predicted cognate CEs, enabled us to test if a given CE was enriched in co-bound versus singly-bound genomic regions. Strikingly, majority of these CEs were significantly enriched in either cobound (red dots) or singly bound (blue dots) regions (Figure 4E, F). Exemplars of such CEs are shown in Figure 4G. The CEs enriched in co-bound regions likely mediate cooperative DNA binding by their cognate TFs whereas those in singly-bound regions are consistent with an anticooperative mode of DNA interaction. Intriguingly, the CE types that were enriched in singly-bound regions manifested a range of configurations with instances of partially overlapping motifs as well as large spacer lengths, implying that anti-cooperative DNA binding is not necessarily a consequence of steric hindrance (discussed below).

Finally, we sought to validate the functional relevance of CEs. To do this, we analyzed their distributions within a large set of active enhancers (~12,000) in murine B cells that had been delineated by us with chromatin profiling and STARR-seq coupled with Hi-C(*22*). Subsets of CEs for TFs that regulate B cell identity and function, including E2A, PU.1, Pou2f1, Mef2C, Ikaros and Blimp1, were preferentially enriched in STARR-seq positive regions (active enhancers) compared with STARR-seq negative regions (inactive OCRs) (Figure 4H, red dots). Notably, distinctive CEs for these TFs were also reciprocally enriched in STARR-seq negative regions (blue dots). Such genomic regulatory regions could reflect poised enhancers or silencers. Nevertheless, the enrichment of ImmCEs, for biologically important B cell TFs, in a large set of functionally validated enhancers testifies to their importance. In our analysis of the B cell cis-regulome, STARR-seq enhancers were connected with their interacting promoters using Hi-C thereby revealing mono-genic (enhancers interacting with single promoters) as well as multi-genic enhancers (enhancer interacting with 8 or more promoters) configurations(*22*). This provided us with an opportunity to test for possible functional distinctions between *digital* and *fuzzy* CEs. Strikingly, *digital* CEs were enriched in mono-genic enhancers whereas their *fuzzy* counterparts were prevalent in multi-genic enhancers (Figure 4I). Furthermore, analysis of the active enhancer landscape of B cells on the basis of recruitment of the transcriptional co-activator p300, revealed *fuzzy* CEs to be enriched in enhancers occupied by p300 (Figure 4J). Notably, we have previously shown that multi-genic enhancers display increased activating histone modifications (H3K27Ac, H3K4me3) with larger numbers of interacting promoters(*22*) and this maybe mediated in part by increased p300 recruitment. Importantly both *digital* and *fuzzy* CEs are enriched in active enhancers but partitioned differentially into mono-genic and multi-genic enhancers, respectively. The stereospecific constraints inherent in *digital* CEs may contribute to the specificity of monogenic enhancers whereas the relaxation of TF motif spacing constraints in *fuzzy* CEs and the increased p300 recruitment, may promote engagement of multiple promoters by multi-genic enhancers.

Our systematic analysis of a large set of OCRs, spanning the chromatin landscape of a multitude of immune and hematopoietic cell types and states, has revealed a diverse set of CEs that are a prevalent second order feature of regulatory genomic sequences. Given that the CEs involve a diverse set of TF motif pairs that can be bound by members of TF families expressed in diverse cell types, we anticipate that these CEs can function in many other mammalian cell contexts. From a historical standpoint a two-tiered organizational structure, comprising of simple and binary TF motifs, of the mammalian regulatory DNA code had been predicted over 30 years ago by molecular genetic dissection of the first mammalian enhancer (SV40)(*29*). However, this speculative model could not be elaborated on because of lack of high throughput biochemical and genomic methodologies. It is notable that in spite of considerable progress in chromatin profiling and computational genomics, current analyses of regulatory sequences continue to rely on simple TF motifs for decoding their structures and testing predictions with perturbation experiments. Each CE in our catalog generates strong biochemical and gene regulatory predictions that can be experimentally tested with molecular, genomic and biological experimentation. These extensive experimental analyses will ultimately require consortium-wide efforts such as those organized by ImmGen or ENCODE. Nonetheless, the CEs revealed by CEseek represent a new major resource for TF motif and gene regulatory network inference and open up a vast but underappreciated regulatory genomic landscape for exploration.

Our conclusion that CEs represent molecularly and functionally relevant TF motif configurations critically relies on three complementary approaches for its validation. These include matching of CEs with sequences in a large CAP-SELEX dataset that involves affinity-based selection of synthetic sequences recognized by cooperatively binding TF pairs(*19*). Thus, many CEs clearly have a biochemical underpinning. Furthermore, a large number of CEs are enriched in TF ChIP-seq datasets with a subset reflected in co-bound DNA regions of the predicted TF pair. Finally, CEs are also shown to be enriched in active enhancers within a STARR-seq dataset. In spite, of this extensive validation we recognize that many CEs within our catalog of ~20,000 remain to be molecularly and functionally assessed. This could be achieved by high throughput approaches such as protein-binding microarrays(*30, 31*) along with massively parallel reporter assays(*32*) complemented with comprehensive profiling of binding regions of hundreds of TFs in diverse cellular contexts. A limitation of our approach is that it has focused on a search space of regulatory sequences that are primarily operative in immune and hematopoietic cells. Although it is likely that many of the discovered ImmCEs will be biologically relevant in other cell contexts, nevertheless we anticipate that the CE universe will expand significantly by extension of our approach to additional mammalian cell types.

A notable finding is that CEs are reflected by *digital* and *fuzzy* configurations of their simple TF motifs that comprise them. *Digital* CEs have been classically defined by exemplar composite elements that are constituted by stereospecific juxta positioning of two TF motifs that promote cooperative DNA binding of the TFs. As expected, many predicted *digital* CEs can be cooperatively recognized by TF pairs (CAP-SELEX dataset) and are enriched in co-bound genomic regions of the predicted pairs of TFs (ENCODE ChIP-seq datsets)(*28*). We note that in some *digital* CEs, particularly with partial overlap of the simple TF motifs there can be a significant deviation of one or both of the individual sequence motifs. This is very likely a consequence of new modes of DNA recognition by TF pairs as a consequence of TF-TF interactions or DNA allostery as has been documented in a few instances(*33, 34*). Intriguingly, many *digital* CEs are depleted in co-bound genomic regions of their cognate TFs, instead being enriched in singly bound regions. This property strongly suggests an anti-cooperative mode of DNA recognition of such CEs by cognate TF pairs as has been documented in several cases by us and others(*26, 27*). Our analysis of *digital* CEs reveals a highly diverse set of TF motif pairs which strongly suggests that structurally distinctive TF families can participate in cooperative or anti-cooperative modes of DNA binding. However, it should be noted that CEs could also be used to discriminate the actions of TFs that belong to the same family by selectively enabling cooperative or anti-cooperative binding. This has been seen to be the case for the EICE motif in specifically enabling cooperative binding by PU.1 or Spi-B (Ets family) and IRF4 or IRF8 (IRF family) but not by other members of the Ets and IRF family members(*9, 14*) as well as for an Ets-AP1 CE (Supplementary Figure S6)(*27*). In contrast with their *digital* counterparts *fuzzy* CEs are notable in lacking stereospecific constraints of their parent TF motifs up to spacer lengths of 10 bp. Despite the extreme variability of their configurations, *fuzzy* CEs are notable in that they are comprised by structurally less diverse TF motif pairs. This is a paradoxical finding that will require detailed molecular studies of TF interactions with *fuzzy* CEs to adequately resolve. The molecular basis of cooperative DNA binding of TF pairs to very heterogeneous CE configurations remain to be elucidated. A plausible mechanism would invoke a flexible molecular scaffold such as the RNA Pol II mediator complex or the co-activators P300/CBP that preferentially promote interactions between the two DNA bound TFs(*35*). Consistent with such a model, we show *fuzzy* CEs to be selectively enriched in genomic regions bound by p300. From a functional standpoint *fuzzy* CEs are strongly enriched in multi-genic enhancers suggesting that they may facilitate the interaction of each such enhancer with multiple cognate promoters via stronger interactions with the forementioned molecular scaffolds such as p300.

Recent studies have used different computational inference models to analyze the syntax of the genomic regulatory code (*36, 37*). One of these has explored the possibilities of altering consensus motifs for the TFs of the same family based on the resulting shape of motifs(*37*). Another study has investigated the possibility of soft-syntax for the regulatory code using helical distance to expand the spectrum of binding preferences(*36*). These studies can be considered as exemplars of top down approaches. We have used a bottom up approach informed by DNA binding properties of TF pairs to systematically analyze a secondary layer of organization within the genomic regulatory code. It remains to be determined if our approach can be extended to higher levels of organization. Both types of approaches, unbiased deep machine learning as well as hierarchical analyses based on structural properties of DNA-protein interactions will be necessary to thoroughly delineate the regulatory code. The latter code though analogous with the genetic code is far more complex and still remains a formidable challenge to solve in the era of genomics.

## Acknowledgements

We thank Fangping Mu and the Center for Research Computing (CRC) at University of Pittsburgh for assistance with and access to the computational resources. We are extremely appreciative of the ImmGen and ENCODE consortium efforts without which this work would not have been possible. We are grateful to Neha Cheemalavagu, Syed Rahman and Jingyu Fan for testing CEseek and providing useful feedback with the end user experience. We thank Jishnu Das, Lee Grimes, Nathan Salomonis, Rachel Gottschalk, Takis Benos, Jian Ma, Mark Shlomchik and Singh Lab members (Jingyu Fan and Louis Lau) for their critical and constructive comments.

## Author Contributions

VKC and HS conceived the general conceptual framework and various types of analyses to be performed. VKC developed the CEseek pipeline and performed all computational analyses of the datasets. HS and VKC wrote the manuscript together.

## Data and Material availability

All datasets used for computational analyses are cited with their primary sources. CEseek is available at https://github.com/viren-v/CEseek

## Supplementary Figure Legends

**Supplementary Figure S1.**
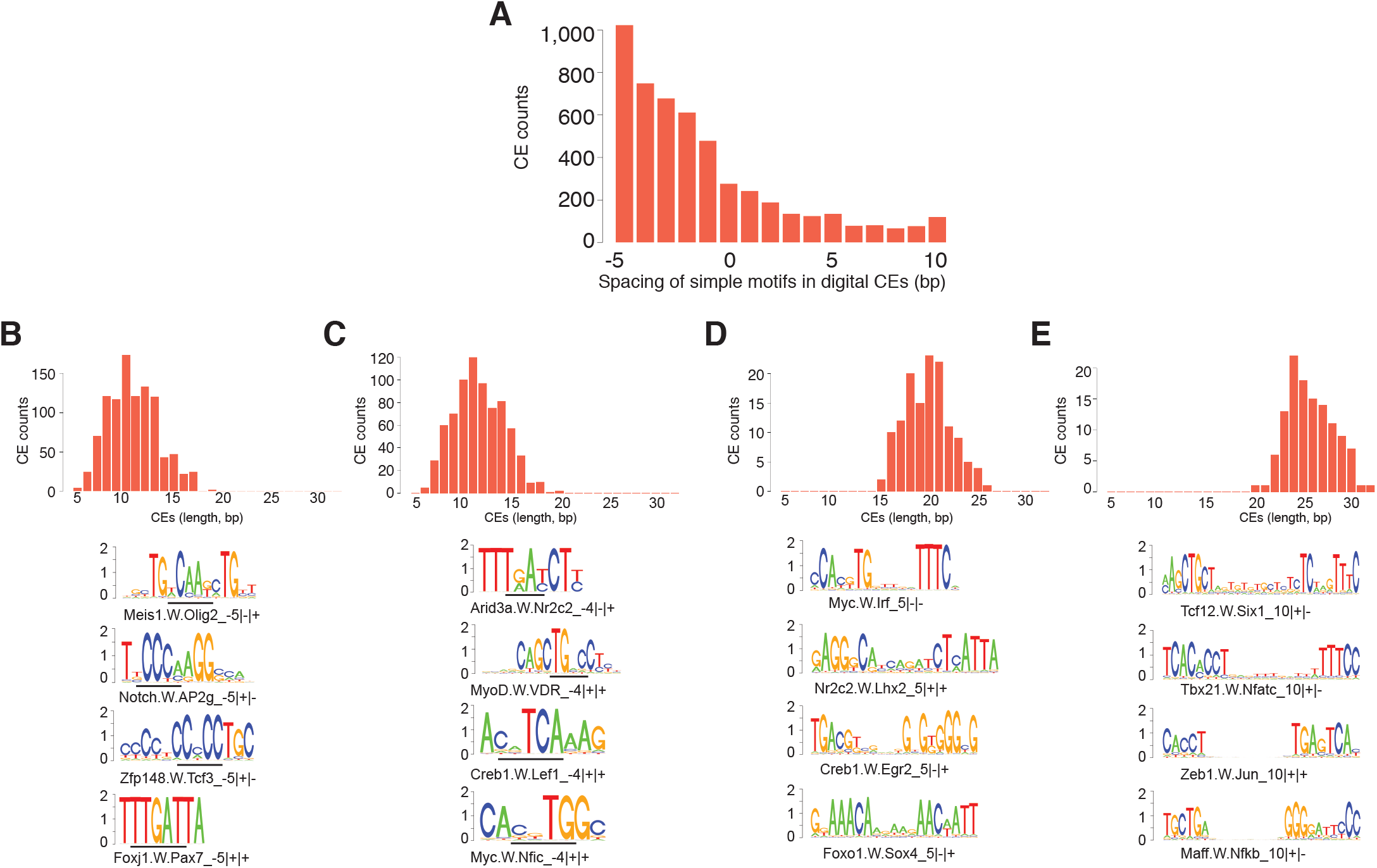
Features of digital CEs. (A) Bar plot displays counts of digital CEs for indicated spacer lengths of partner TF motifs. (B, **C**) Distribution of CE counts for overlapping digital CEs with −5, −4, spacer configurations, respectively. Exemplary CEs are shown below each distribution as motif logos. Bars highlight the overlapping positions of simple TF motifs comprising the CEs. (**D, E**) Distribution of CE counts for non-overlapping digital CEs with 5bp and 10bp spacer configurations, respectively. Exemplary CEs are shown below each distribution as motif logos.

**Supplementary Figure S2.**
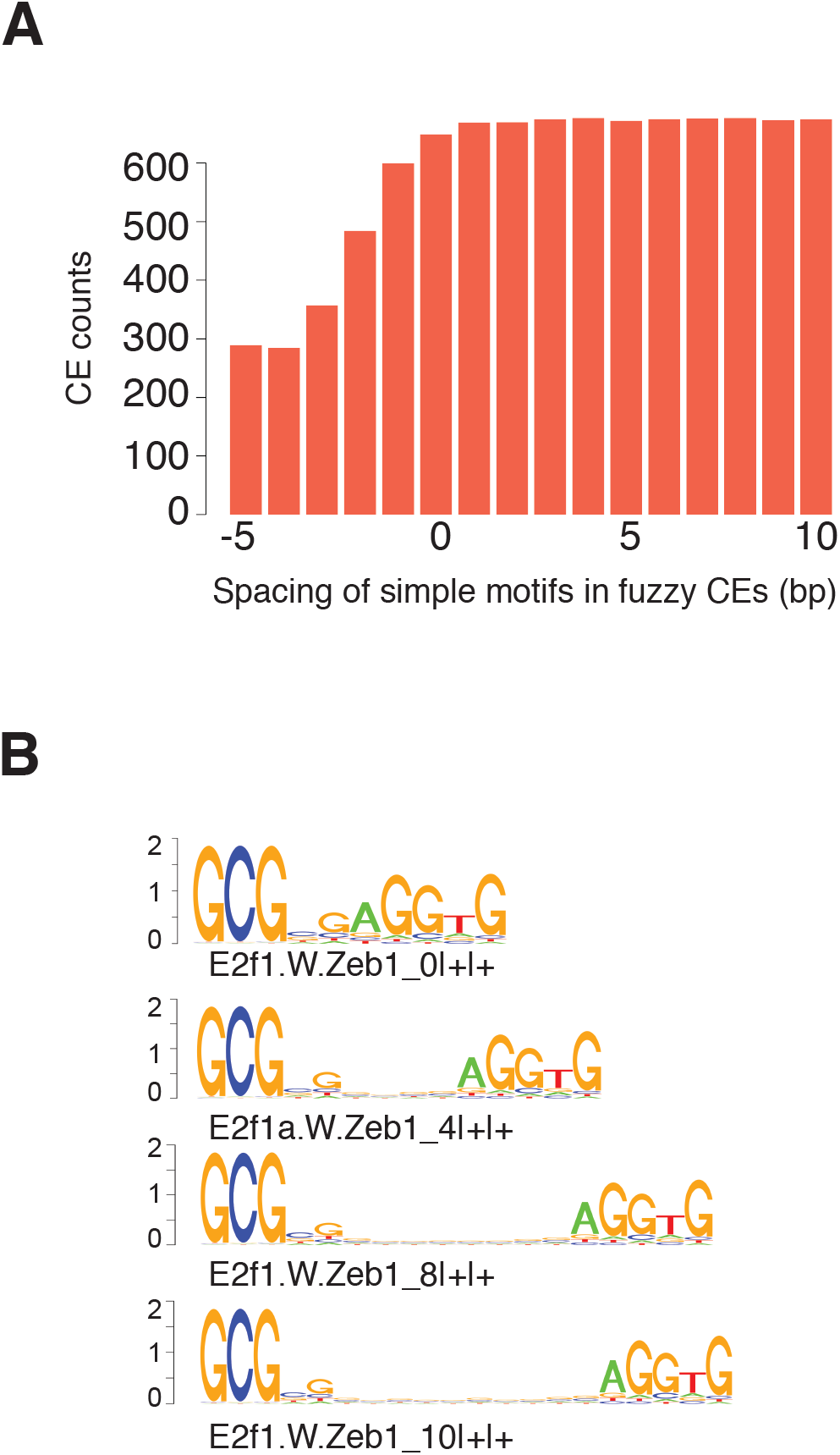
Features of fuzzy CEs. (A) Bar plot displays counts of fuzzy CEs for indicated spacer lengths of partner TF motifs. (B) Exemplary fuzzy CE comprised by E2F and Zeb motifs with 4 out of 53 statistically enriched configurations displayed as motif logos (spacer lengths of 0, 4, 8 and 10bp).

**Supplementary Figure S3.**
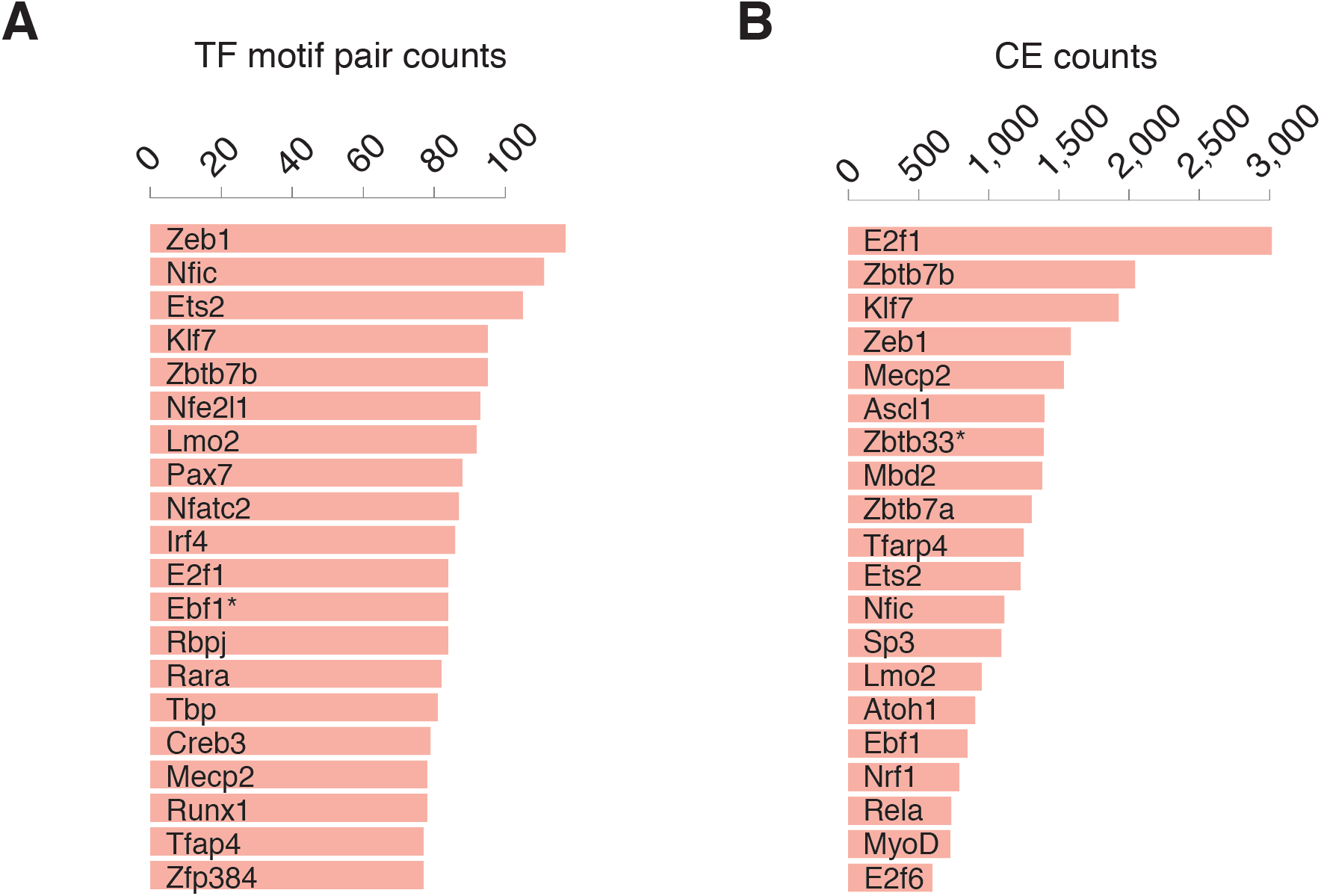
Dominant TF motifs in CE types versus CE configurations. (A) Bar plot shows the top 20 TF motifs and their counts of distinct motif pairs, comprised by the pairing of indicated motif with different partner motifs (CE types). (B) Bar plot shows the top 20 TF motifs with counts of total CEs that include the indicated motif with different partner motifs and all of their statistically enriched configurations. Asterisks signify monomeric motifs for the indicated TFs comprise the relevant CEs.

**Supplementary Figure S4.**
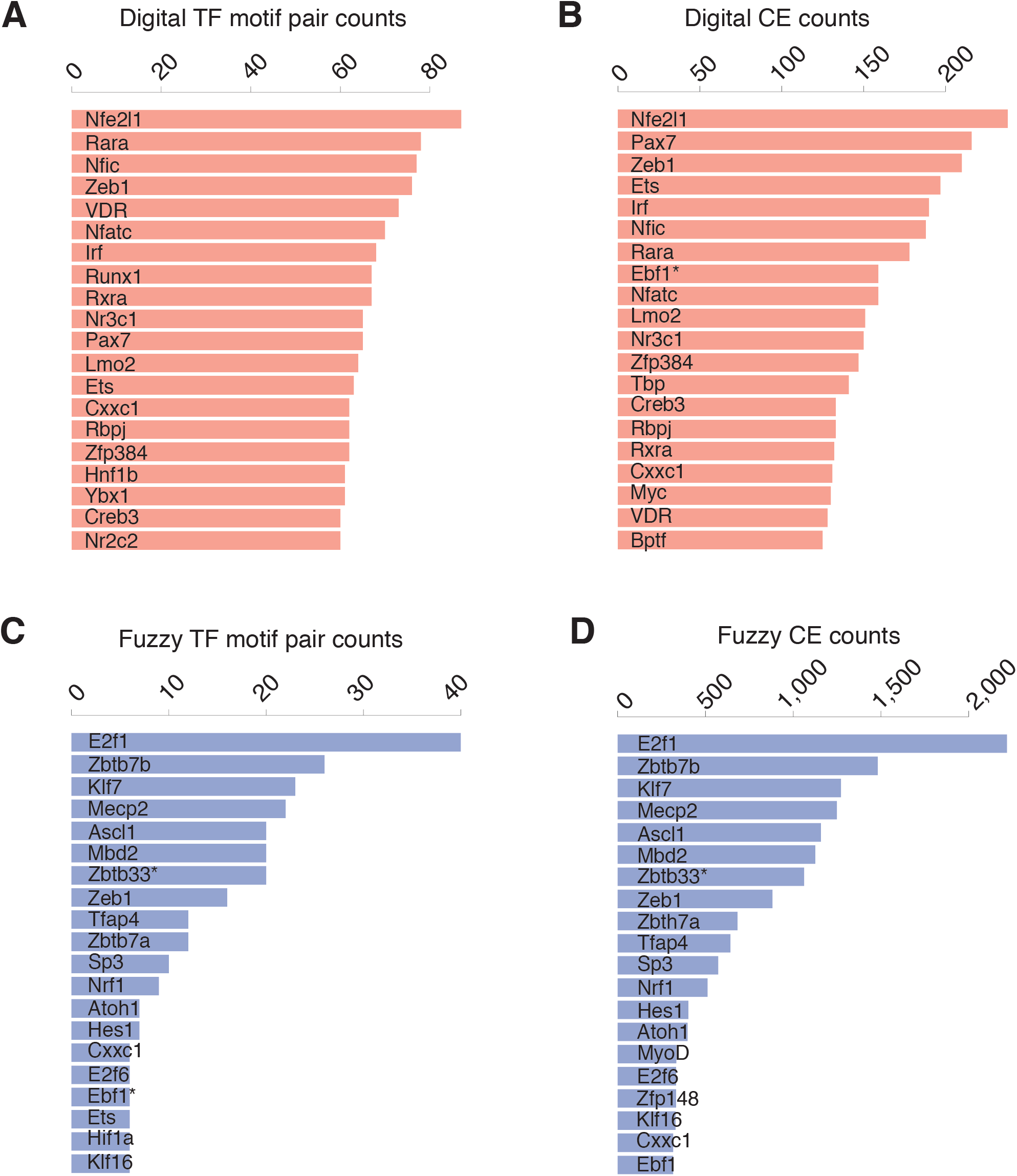
Dominant TF motifs in digital versus fuzzy CEs. (A, B) Bar plots show the top 20 TF motifs for digital CE types and configurations as in Supplementary Figure S3. (C, D). Bar plots show the top 20 TF motifs for fuzzy CE types and configurations as in Supplementary Figure S3. Asterisks signify monomeric motifs for the indicated TFs comprise the relevant CEs.

**Supplementary Figure S5.**
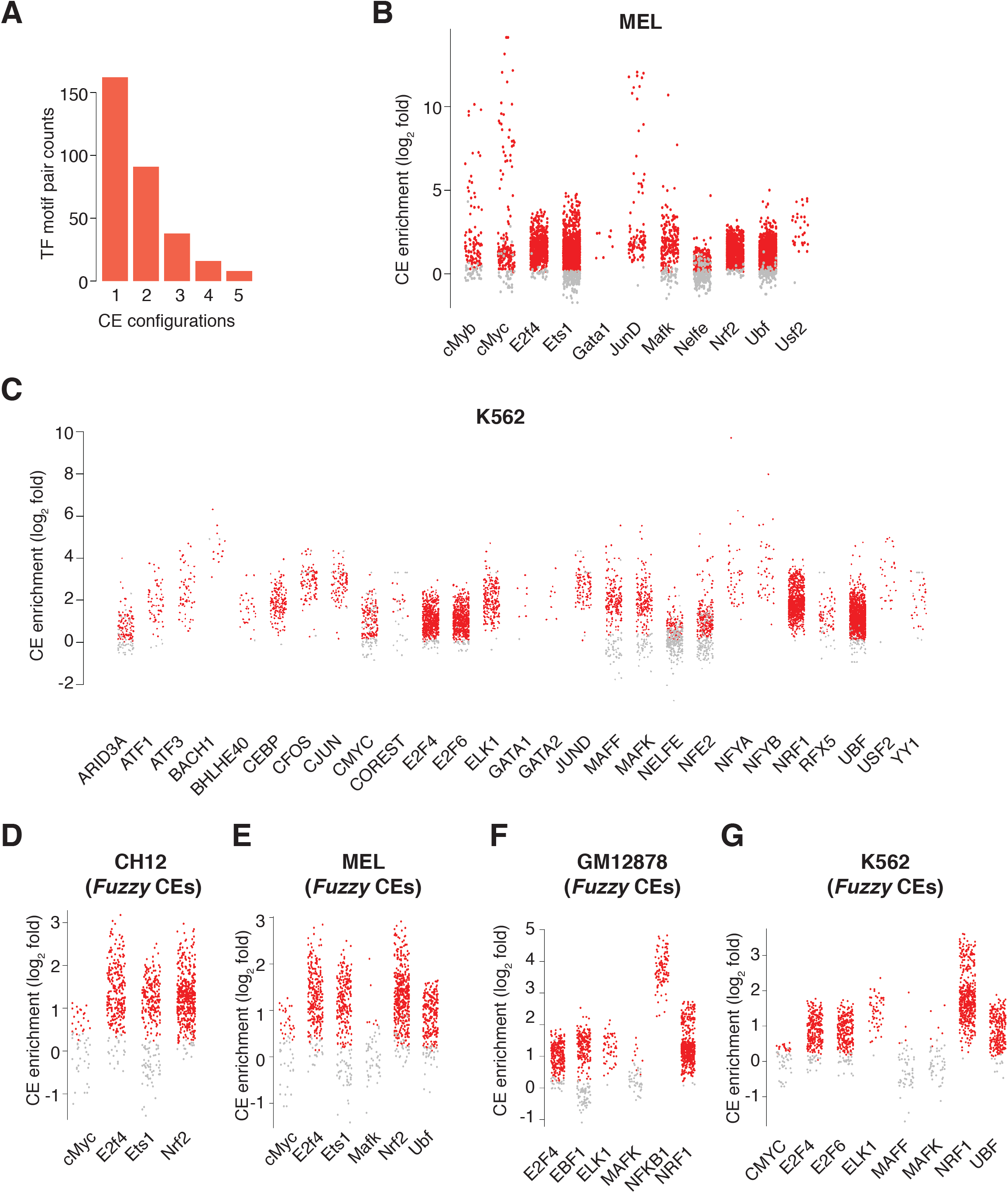
CEs are enriched in the genomic binding regions of a diverse set of murine and human TFs. (A) Bar plot displays counts of CEs identified by Cap-SELEX (n=456) as a function of their numbers of configurations (1 to 5). (B) Enrichment of CEs for the indicated murine TFs in their ChIP-seq peaks in the murine MEL erythroleukemia cell line. Background set of sequences (10x) was constituted by scrambling the test set. Red dots indicate CEs with an enrichment significance of logP < −20. Grey dots are CEs that do not satisfy the significance criterion. (C) Enrichment of CEs for the indicated human TFs in their ChIP-seq peaks in the human K562 erythroleukemia cell line. Analysis and color coding of CEs is same as in panel B. (D) Enrichment of fuzzy CEs in ChIP-seq datasets for indicated TFs in CH12 cell line. (E) Enrichment of fuzzy CEs in ChIP-seq datasets for indicated TFs in in MEL cell line. (F) Enrichment of fuzzy CEs in ChIP-seq datasets for indicated TFs in GM12878 cell line. (G) Enrichment of fuzzy CEs in ChIP-seq datasets for indicated TFs in K562 cell line. Analysis and color coding of CEs is same as in panel B. The Cap-SELEX and ChIP-seq datasets are same as in Fig. 4.

**Supplementary Figure S6.**
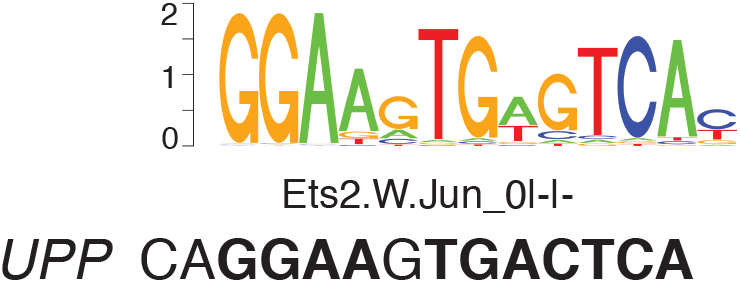
An Ets-AP1 CE shown to promote cooperative as well as anti-cooperative DNA binding by Ets and AP1 family members. The motif logo for an Ets-AP1 CE delineated by CEseek is displayed. The sequence in black font is derived from the promoter of the *UPP* gene and shown to promote cooperative binding by ERG and AP1 (FOS-JUN dimer)(*27*). Conversely, binding of the CE by the Ets family member EHF is inhibited by AP1 (FOS-JUN dimer).

## Supplementary Tables Description

Supplementary Table 1 Simple TF motif library (PPMs)

Supplementary Table 2 ImmCEs predicted by CEseek scan of ImmGen ATAC-seq datasets (PPMs)

Supplementary Table 3 CEs with high similarity to parent simple motifs (similarity score >0.9)

Supplementary Table 4 Enrichment statistics of CEs using a 5-fold scrambled set as background

Supplementary Table 5 Catalog of Digitial CEs (PPMs)

Supplementary Table 6 Catalog of Fuzzy CEs (PPMs)

Supplementary Table 7 TF motif pair counts for indicated simple TF motifs (All CEs, Digital CEs and Fuzzy CEs)

Supplementary Table 8 CE counts for indicated simple TF motifs (All CEs, Digital CEs and Fuzzy CEs)

Supplementary Table 9 Enrichment statistics of CEs in cognate ChIPseq datasets from CH12, GM12878, MEL and K562 cells

Supplementary Table 10 Enrichment statistics of CEs in co-bound regions of cognate TF pairs in ChIPseq datasets from CH12, MEL, GM12878 and K562 cells

Supplementary Table 11 Enrichment statistics of CEs for indicated TFs in B cell STARR-seq enhancers

Supplementary Table 12 Enrichment of digital and fuzzy CEs in monogenic vs multigenic B cell STARR-seq enhancers

Supplementary Table 13 Enrichment of digital and fuzzy CEs in P300 bound vs not-bound B cell STARR-seq enhancers

## MATERIALS AND METHODS

### CEseek pipeline

CEseek is written and implemented in R (*1*). It uses R packages GenomicRanges (*2*) for genomic interval operations and Biostrings for handling DNA sequences and scanning TF motifs using position probability matrix libraries (PPMs). The inputs are DNA sequences as DNAStringSet objects and PPMs for TFs as an R list object. Biostrings function matchPWM is employed with a user specified stringency (% match) for calling hits. TF motif hits described herein were based on a match stringency of 80%. The TF motif hits are scanned for in a given sequence in a pairwise manner, for all four orientations and user specified spacing intervals. For the binary configurations detailed herein, −5 to +10 bp end to end spacing between a given pair of TF motifs was employed. Hits for each binary configuration are extracted using the relevant ends of the two TF motifs including the spacer. These binary TF motif hits are stored in a list structure with an identifier of the sequence from which they are extracted. Herein, each sequence identifier is linked to its genomic coordinates and accessibility defined by ATAC-seq or enhancer activity delineated by STARR-seq in a particular immune cell type or state. The aligned binary TF motif hits are used to generate PPMs of the corresponding CEs. The Fisher’s exact test is used to determine if a particular CE is statistically enriched in the test sequences using a suitable background set of sequences. CEseek is available at https://github.com/viren-v/CEseek

### CEseek scan of B cell enhancers

The proof-of-concept CEseek was performed on 11,809 STARR-seq positive regions identified in activated B cells (*3*) using a 10-fold scrambled set of DNA sequences as background. Input TF motifs (PPMs) represented those for IRF, PU.1, AP1 and NFAT proteins.

### Compilation of Simple TF Motif Library for CEseek scan of ImmGen ATAC-seq datasets

Mouse and human TF motifs were downloaded from cis-BP (*4*). A similarity matrix for all PWMs was generated using Homer function “compareMotifs.pl” with option “-matrix”. For any TF with multiple PWMs a minimal spanning tree, with the PWMs as nodes, was produced using the similarity matrix. The PWM with most connecting edges was selected and the others discarded. In the case of ties, the PWM with highest information content was chosen. In cases where 2 or more TFs had PWMs with similarity score of more than 0.9 then all were represented by a single PWM with the highest information content. Gene symbols of TFs with similar DNA specificities were separated by “:”in the nomenclature of the simple motifs in the final library. For completeness, the cis-BP library was supplemented with curated TF motifs from the Homer vertebrate TF motif library(*5*). These ChIP-seq derived motifs represented core (monomeric) sequences. The final compiled TF motif library was manually checked so as to reduce the number of very similar motifs. The final library is comprised of 153 simple TF motifs (PPMs, Supplementary Table 1). Motif logos can be generated using these PPMs on http://www.benoslab.pitt.edu/stamp/.

### CEseek scan of ImmGen ATAC-seq datasets

Chromatin profiles (ATAC-seq datasets) of 86 hematopoietic and immune cell types generated by the ImmGen consortium were used for the CEseek scan. This catalog of open chromatin regions (OCRs) comprised >500,000 presumptive transcriptional regulatory sequences (*6*). ATAC-seq peak co-ordinates were extended for 90 bp on each side resulting in a 180 bp regulatory region (OCR) corresponding to each peak. This ensured no overlap among the OCRs. DNA sequences corresponding to the OCRs were extracted from the mm10 assembly as DNAStringSet objects in R using BSgenome.Mmusculus.UCSC.mm10 package. Sequence annotations were carried over from the ImmGen resource (*6*).

The ImmGen OCR sequences were scanned with CEseek for all possible TF motif combinations and configurations (−5 to +10 bp spacing) using the simple TF motif library comprised of 153 core PPMs. The scan was performed in batches of 500 OCR sequences on the University of Pittsburgh Center for Research Computing cluster. The CEseek outputs for all batches were merged resulting in approximately 600,000 configurations with at least 1 occurrence in the OCR dataset.

For stastistical enrichment analysis of CEseek configurations, each OCR (ATAC-seq peak) was first assigned to one or more cell types or cell states based on its preferential accessibility (95th percentile) enumerated by signal intensity across all ImmGen cell states. These OCRs are referred to as cell type or cell state-specific regulatory sequences. Fisher’s test was then used to determine if a particular CEseek configuration (total ~600,000) was enriched in a given set of cell type or state-specific OCRs using the rest of ImmGen OCRs as the background set. A p-value of less than 1.0E-20 was used as a cut-off for statistical enrichment, resulting in delineation of ~20,000 CEs. In a final step, PPMs for the qualifying CEs were generated using all corresponding hits of CE in the entire ImmGen OCR dataset (Supplementary Table 2).

An independent test for the prevalence of these CEs in ImmGen OCRs was based on a complementary approach involving a set of scrambled sequences as background. This alternative criterion revealed 16,839 of the CEs to be significantly enriched in the test set with p values of less than 10^-10^ (Supplementary Table 4).

### Minimal spanning tree analysis of ImmGen cell types and states based on CE prevalence

To generate a minimal spanning tree (MST) that relates diverse immune cell types and states in the ImmGen datasets, a matrix was generated using the fractional coverage of cell type/state specific OCRs by CEs in the top 2 quartiles of the CE distribution in Figure 2D. Fractional coverage is determined by enumerating the percentage of cell type/state specific OCRs that are assigned a given CE by CEseek. The matrix was converted into distance object using function “dist” in R. MST was calculated using function MST from package “ape”. Resulting MST object was converted in graph and plotted using package “igraph”.

### CAP-Selex analysis

Position Frequency Matrices (PFMs) of interacting TFs analyzed by CAP-Selex (*7*) were converted into PPMs. This enabled a comparison between CEs and the CAP-Selex motifs using homer function compareMotifs.pl. Highly similar TF motifs in CAP-selex datasets were collapsed by Homer to yield a final set of 455. Best match scores were used to plot the data.

### ChIP-seq analysis of ENCODE datasets

Bam files of ChIP-seq datasets were obtained from UCSC’s ENCODE download portal (http://hgdownload.cse.ucsc.edu/goldenPath/hg19) and then processed using homer as described (*3*). Peaks overlapping with blacklisted regions were discarded. Homer was used for enrichment analysis of CEs in ChIP-seq peaks using a 10-fold scrambled set of sequences generated using homer function “scrambleFasta.pl”. For determining single-bound vs. co-bound regions the peaks were merged for pairs of TFs in R as GRanges object. Function “reduce” was used to convert overlapping regions into contiguous unique regions. Regions overlapping with peak(s) from the corresponding TF-ChIP-seq datasets were considered co-bound, while the remaining regions were designated as singly bound. The co-bound and singly bound regions were then tested for cognate CE enrichment using Homer.

